# FtsW is a peptidoglycan polymerase that is activated by its cognate penicillin-binding protein

**DOI:** 10.1101/358663

**Authors:** Atsushi Taguchi, Michael A. Welsh, Lindsey S. Marmont, Wonsik Lee, Daniel Kahne, Thomas G. Bernhardt, Suzanne Walker

## Abstract

The peptidoglycan cell wall is essential for the survival and shape maintenance ofbacteria.^1^ For decades it was thought that only penicillin-binding proteins (PBPs) effected peptidoglycan synthesis. Recently, it was shown that RodA, a member of the Rod complex involved in side wall peptidoglycan synthesis, acts as a peptidoglycan polymerase.^2–4^ RodA is absent or dispensable in many bacteria that contain a cell wall; however, all of these bacteria have a RodA homologue, FtsW, which is a core member of the divisome complex that is essential for septal cell wall assembly.^5,6^ FtsW was previously proposed flip the peptidoglycan precursor Lipid II to the peripasm,^7,8^ but we report here that FtsW polymerizes Lipid II. We show that FtsW polymerase activity depends on the presence of the class B PBP (bPBP) that it recruits to the septum. We also demonstrate that the polymerase activity of FtsW is required for its function *in vivo*. Our findings establish FtsW as a peptidoglycan polymerase that works with its cognate bPBP to produce septal peptidoglycan during cell division.

---

Peptidoglycan is synthesized from the cell wall precursor Lipid II. This lipid-linked molecule is made inside the cell and then translocated by Lipid II flippases to the extracellular surface of the membrane where it is polymerized and crosslinked (Fig. 1a).^5^ The penicillin-binding proteins (PBPs), a major class of enzymes involved in peptidoglycan synthesis, were identified because they are targets of β-lactam antibiotics such as penicillin, which covalently inactivate their crosslinking domains. The PBPs comprise several types of enzymes, of which the most important for peptidoglycan synthesis are the class A PBPs (aPBPs) and the class B PBPs (bPBPs).^9^ The aPBPs contain a peptidoglycan glycosyltransferase (PGT) domain that polymerizes Lipid II and a transpeptidase (TP) domain that crosslinks the resulting glycan strands (Fig. 1a). The bPBPs consist of a TP domain and a domain of unknown function. Some bacteria also contain monofunctional glycosyltransferases (MGTs) that have homology to the PGT domains of the aPBPs, and these proteins can also polymerize Lipid II. It is assumed that the glycan strands produced by MGTs are crosslinked by PBPs.

**Figure 1:**
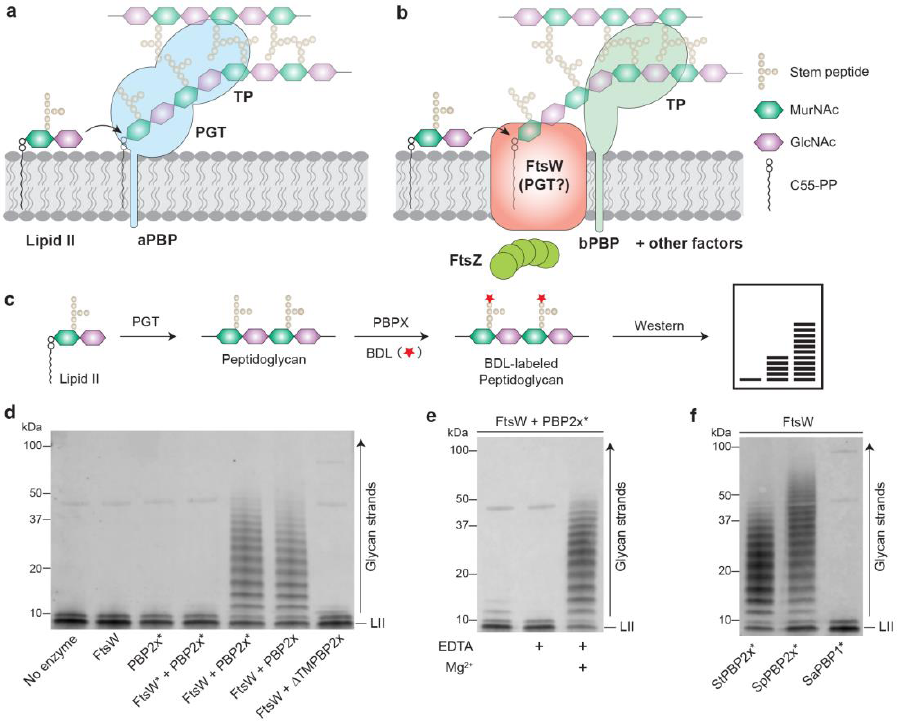
FtsW is a peptidoglycan synthase. **a**, Peptidoglycan synthesis by the bifunctional class A PBPs (aPBPs). b, Septal peptidoglycan synthesis is directed by the divisome. FtsW may have peptidoglycan polymerase activity like its homologue RodA from the Rod system.^2^ c, Schematic of the PAGE assay used to detect peptidoglycan generated *in vitro*. **d**, *S. thermophilus* FtsW polymerizes Lipid II in the presence of its cognate bPBP, PBP2x. **e**, FtsW requires divalent cations for polymerase activity. **f**, *S. thermophilus* FtsW can be activated by a near-cognate bPBP from *S. pneumoniae*. Asterisk (*) indicates the catalytically inactive variant.

The β-lactam antibiotics are the most clinically and commercially successful class of antibiotics in history. These drugs continue to be widely used for treating infections, but the emergence of β-lactam resistance in major human pathogens makes the identification of new antibacterial targets a high priority. Proteins that are essential in all bacteria, such as those belonging to the SEDS (shape, elongation, division, and sporulation) family, are particularly appealing therapeutic targets. Phylogenetic analysis has shown that the SEDS proteins RodA and FtsW are more widely conserved than the aPBPs.2 RodA is essential in rod-shaped organisms where it serves as a key component of the Rod complex (elongasome) that makes side wall peptidoglycan.^9^ FtsW, on the other hand, is required for the function of the division machinery (divisome) and is essential in most bacteria regardless of their shape.^9^

SEDS proteins have been found to form subcomplexes with bPBPs.^10,11^ Therefore, the recent discovery that RodA has peptidoglycan polymerase activity has led to the proposal that SEDS-bPBP complexes form two-component peptidoglycan synthases with polymerase and crosslinking activities on separate polypeptides (Fig. 1b).^2–4^ Accordingly, mutations predicted to disrupt RodA interactions with its cognate bPBP (PBP2) cause the Rod system to malfunction *invivo*.^12^ However, polymerase activity has thus far only been demonstrated for RodA from *Bacillus subtilis*.^2^ Whether other SEDS proteins like FtsW also function as peptidoglycan polymerases remains unclear as does the effect of association with bPBPs on this activity. We therefore expressed and purified FtsW alone or in complex with its cognate bPBP from several organisms to investigate their ability to function as peptidoglycan synthases.

We first expressed and purified FtsW orthologs from the Gram-positive bacteria *Streptococcus thermophilus* (*St*FtsW) and *Staphylococcus aureus* (*Sa*FtsW) (Supplementary Fig. 1), but failed to detect polymerase activity (Fig. 1c) when the proteins were incubated with their native Lipid II substrate (Fig. 1d and Supplementary Fig. 2). Strikingly, addition of the cognate bPBP to the FtsW reactions resulted in the production of peptidoglycan oligomers consistent with the stimulation of peptidoglycan polymerase activity (Fig. 1d and Supplementary Fig. 2). This activity was abolished when an invariant aspartate in or near the presumed active site in FtsW was mutated to alanine, but not when the TP active site of the bPBP was inactivated (Fig. 1d and Supplementary Fig. 2). Furthermore, like RodA, the activity of the *St*FtsW in the presence of its cognate bPBP (*St*PBP2x) was insensitive to the phosphoglycolipid antibiotic moenomycin that inhibits aPBPs and MGTs (Supplementary Fig. 3).^2^ Another distinction between the activity of FtsW and that of aPBPs and MGTs is a divalent cation requirement for FtsW polymerase activity. When EDTA was added to the buffer, *St*FtsW polymerase activity was abolished, but adding excess Mg^2+^, Ca^2+^, or Mn^2+^ to the EDTA-containing buffer restored polymerization (Fig. 1e and Supplementary Fig. 4). We therefore conclude that FtsW is a peptidoglycan polymerase that requires divalent cations and the presence of a bPBP for activity.

Studies in *E. coli* have demonstrated that the transmembrane (TM) domain of PBP3 is sufficient for its septal localization, and evolutionary coupling analyses have suggested that the TM domain of bPBPs interact with SEDS proteins.^12–15^ To test whether the TM domain of a cognate bPBP is required for the PGT activity of FtsW, we removed the TM domain from *St*PBP2x and tested the ability of the ΔTM variant to activate *St*FtsW. No peptidoglycan product was detected, showing that the TM helix is crucial for promoting FtsW polymerase activity (Fig. 1d). To assess the specificity of the polymerase promoting activity, *St*FtsW and *Sa*FtsW reactions were performed in the presence of divisome bPBPs from three different species (Fig. 1f and Supplementary Fig. 5). *Sa*FtsW was only able to polymerize Lipid II when combined with its native bPBP, PBP1 (*Sa*PBP1), but *St*FtsW was able to polymerize Lipid II when combined either with its cognate bPBP, *St*PBP2x, or with the divisome bPBP from *S. pneumoniae* (*Sp*PBP2x), which is 50% identical to its *S. thermophilus* counterpart. *St*FtsW showed no activity in the presence of *Sa*PBP1. Therefore, FtsW polymerase activity requires its specific cognate or near-cognate bPBP, and the TM helix of the bPBP likely forms part of the protein-protein interface between the PGT and TP enzymes.

To determine if FtsW forms a stable peptidoglycan synthase complex with its cognate bPBP in both Gram-positive and Gram-negative bacteria, we co-expressed *Sa*FtsW with its partner *Sa*PBP1, and FtsW from *Pseudomonas aeruginosa* (*Pa*FtsW) along with its cognate bPBP, *Pa*PBP3. In both cases, the bPBP co-purified with the affinity-tagged FtsW, and the complexes possessed PGT activity (Figs. 2a, 2c and Supplementary Fig. 6). As with the individually purified components, the activity of the complexes was inactivated by substitution of the invariant aspartate in or near the presumed active site of FtsW (Figs. 2a, 2c and Supplementary Fig. 6). To assess the *in vivo* relevance of the detected FtsW polymerase activity, we compared the effect of overproducing either wild-type FtsW or inactive FtsW variants (designated FtsW*) in their native host organism. Cell growth and division remained relatively normal upon expression of wild-type FtsW (Fig. 2 and Supplementary Figs. 7 and 8). However, production of *Sa*FtsW* and *Pa*FtsW* inhibited growth of both *S. aureus* and *P.aeruginosa,* respectively (Supplementary Fig. 7). Moreover, *Sa*FtsW* caused septal abnormalities in *S. aureus*, and *Pa*FtsW* expression induced cell filamentation in *P. aeruginosa* (Figs. 2b, 2d, Supplementary Figs. 8 and 9). The observed dominant-negative activity of the FtsW* proteins in their host organism most likely results from the defective proteins saturating FtsW binding sites in the divisome complex to disrupt its function.

**Figure 2:**
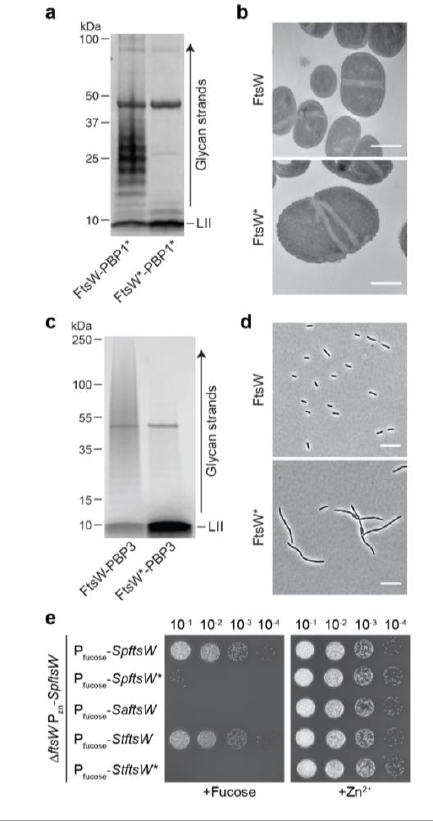
The PGT activity of FtsW is essential for cell division. **a**, *In vitro* polymerization of Lipid II by co-purified *S. aureus* FtsW-PBP1* complexes. **b**, Electron microscopy images of *S. aureus* cells overexpressing the wild-type FtsW or FtsW*. Scale bar = 500 nm. **c**, *In vitro* polymerization of Lipid II by co-purified *P. aeruginosa* FtsW-PBP3 complexes. **d**, Phase contrast images of *P. aeruginosa* cells overexpressing the wild-type FtsW or FtsW*. Scale bar = 10 µm. **e**, Depletion of FtsW in *S. pneumoniae* can be rescued by the expression of *SpftsW* or *StftsW*.

To further investigate the essentiality of FtsW polymerase activity, an FtsW depletion strain of *S. pneumoniae* was constructed. Expression of a second copy of *SpftsW* or *StftsW* rescued growth upon *Sp*FtsW depletion (Fig. 2e). However, neither of the corresponding *ftsW*\* alleles or *SaftsW* could prevent the lethal *Sp*FtsW depletion phenotype (Fig. 2e). Taken together, the genetic and biochemical results indicate that stable FtsW-bPBP complexes are formed in diverse bacteria and possess peptidoglycan polymerase activity that is required for growth and division.

To evaluate whether FtsW had a reciprocal stimulatory effect on bPBP activity, we used an LC-MS based assay to detect peptidoglycan crosslinks. Peptidoglycan synthesized *in vitro* was digested with mutanolysin (Fig. 3a).^16,17^ These digestions typically produce three readily detectable muropeptide species: disaccharide (monomer) units with an attached pentapeptide (product A), monomer units with an attached tetrapeptide resulting from hydrolysis of the terminal D-alanine of the pentapeptide (product B), and two disaccharide units crosslinked by TP activity (dimer) (product C) (Fig. 3a). When *St*PBP2x was incubated with *St*FtsW and Lipid II, we observed an LC-MS peak in the mutanolysin digestions corresponding to the dimeric product (Fig. 2b and Supplementary Fig. 10). Catalytic inactivation of *St*PBP2x led to the disappearance of this peak, confirming that it resulted from TP activity of this enzyme. We observed the same dimeric species when *St*PBP2x was combined with an MGT from *S. aureus* (SgtB) or with an aPBP containing an inactivating TP mutation, *S. pneumoniae* PBP1a*. Because *St*PBP2x can crosslink peptidoglycan strands produced by these other Lipid II polymerases, we conclude that its activity is not dependent on its cognate FtsW partner.

**Figure 3:**
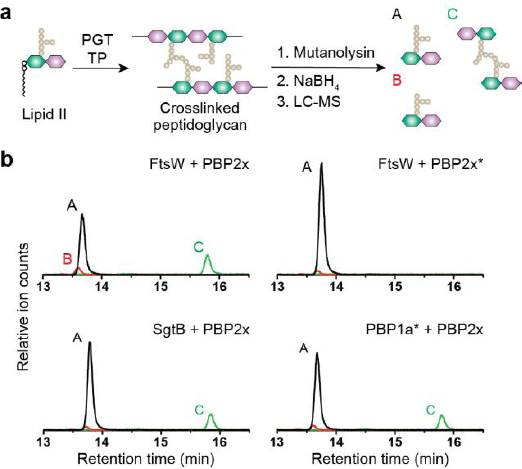
*S. thermophilus* PBP2x does not require FtsW for crosslinking peptidoglycan. **a** Schematic showing detection of crosslinked muropeptide species by LC-MS. Linear peptidoglycan was generated using *S. thermophilus* FtsW, *S. aureus* SgtB or TP inactive *S. pneumoniae* PBP1a* in the presence of *S. thermophilus* PBP2x. Crosslinking is detected by the appearance of peak C, a crosslinked muropeptide. **b,** LC-MS extracted ion chromatograms showing the products of crosslinking reactions with *S. thermophilus* PBP2x. Asterisk (*) denotes the inactivated TP variant.

Our studies have conclusively established that FtsW from several organisms function as a peptidoglycan polymerase and that this activity requires divalent cations and complex formation with a cognate bPBP. The most likely role of the cation is to anchor the diphosphate of the Lipid II substrate and help neutralize its negative charge, but other functions are possible; structural information is needed to better understand the metal requirement. Similarly, how the bPBP partners promote FtsW polymerase activity remains to be determined. An attractive possibility is that the bPBP promotes the formation of an active conformation of FtsW in the complex. This bPBP requirement was not shared by *B. subtilis* RodA as it polymerized peptidoglycan in the absence of its cognate bPBP.^2^ However, this activity was weak and may similarly be stimulated by complex formation.

Recent genetic and biochemical results from our groups also suggest that the polymerase activity of SEDS proteins requires an activation event beyond complexation with a bPBP partner *in vivo*.^18^ Amino acid substitutions in the pedestal domain of PBP2 in *E. coli* were found to stimulate RodA activity *in vivo* and *in vitro*, and these PBP2 variants bypass the requirement for MreC and other components of the shape determining system.^18^ Similar changes in FtsW and PBP3 from *E. coli* and *Caulobacter cresentus* were found to overcome the action of division inhibitors and thus are thought to activate the divisome.^19–21^ It therefore appears that the synthase activity of SEDS-bPBP complexes from both the Rod system and the divisome may be controlled by similar mechanisms within their respective machineries. The development of peptidoglycan polymerase assays for both FtsW and RodA now sets the stage for the molecular details of this regulation to be uncovered.

Another important outstanding question relates to the division of labor between FtsW-bPBP complexes and aPBPs during septal peptidoglycan biogenesis. In *Bacillus subtilis*, cells remain viable in the absence of aPBPs, suggesting that the FtsW-bPBP complex plays a major, possibly sufficient, role in this process.^22^ Septal peptidoglycan synthesis in *B. subtilis* is directly coupled to treadmilling of FtsZ, which colocalizes with PBP2b, the partner of FtsW.^23^ However, most bacterial species require at least one aPBP for viability. In *S. aureus*, the essential aPBP, PBP2, is recruited to the septum and is thought to be crucial for cell division.^24^ Contrasting with *B. subtilis*, the initial rate of septal peptidoglycan synthesis in *S. aureus* is slow and dependent on FtsZ treadmilling, but later in cytokinesis, peptidoglycan synthesis is independent of FtsZ treadmilling.^25^ Thus, FtsW may serve as the major polymerase during treadmilling-dependent cell division, with PBP2 largely responsible for the subsequent treadmilling-independent phase of peptidoglycan synthesis.

Irrespective of their relative roles, both SEDS-bPBP complexes and the aPBPs are clearly critical for proper assembly of the peptidoglycan layer. Further studies of their activity and regulation will therefore pave the way for the discovery of molecules that inhibit their function for future antibiotic development.

## Methods

### Materials

Unless otherwise indicated, all chemicals and reagents were purchased from Sigma-Aldrich. Restriction enzymes were purchased from New England Biolabs. Oligonucleotide primers were purchased from Integrated DNA Technologies. Culture media were purchased from Beckton Dickinson. *S. thermophilus* LMG 18311 genomic DNA (gDNA) was purchased from ATCC (ATCC BAA-250D-5). Biotin-D-lysine (BDL) was prepared by de-protecting Fmocbiotin-D-lysine (Bachem).^17^ *S. aureus* Lipid II and *E. faecalis* Lipid II were isolated from cells as described previously.^16,26^ *S. aureus* SgtB, SgtB^Y181D^, PBP4 and *E. faecalis* PBPX were expressed and purified as previously reported.^16,17,27^

### Bacterial strains, plasmids, oligonucleotide primers and culture conditions

*E. coli* strains were grown with shaking at 37 ºC in lysogeny broth (LB), Terrific Broth (TB, Teknova), or on agarized LB plates. *S. aureus* strains were grown with shaking at 30 °C or 37 °C in Tryptic Soy Broth (TSB) or on agarized plates. *P. aeruginosa* strains were grown with shaking at 30 °C in LB, M9 broth containing 0.2% casamino acids and 0.2% glucose, or on agarized LB plates. *S. pneumoniae* strains were grown statically in Todd Hewitt Broth containing 0.5% yeast extract (THY) at 37 °C in an atmosphere containing 5% CO_2_. When growth on solid media was required, *S. pneumoniae* strains were grown on pre-poured Trypticase Soy Agar with 5% Sheep Blood (TSAII 5%SB) plates with a 5 mL overlay of 1% nutrient broth (NB) agar or TSA plates containing 5% defibrinated sheep blood with appropriate additives. The following concentration of antibiotics were used: ampicillin, 50 µg/mL; carbenicillin, 50 µg/mL; chloramphenicol, 25 µg/mL, erythromycin, 0.2 µg/mL (*S. pneumoniae*) or 5 µg/mL (*S. aureus*); gentamicin, 30 µg/mL; kanamycin, 50 µg/mL (*E. coli*) or 250 µg/mL (*S. pneumoniae*); tetracycline, 0.2 µg/mL. The bacterial strains, plasmids and oligonucleotide primers used in this study are summarized in Supplementary Table 1–3. Protocols for strain and plasmid construction can be found in the Supplementary Methods.

### Protein expression: general procedure

For expression of *Staphylococcus* and *Streptococcus* FtsW and PBPs, *E. coli* C43(DE3) containing the expression plasmid was grown in 1 L TB supplemented with kanamycin or carbenicillin at 37 °C with shaking until OD_600_ was 0.7-0.8. The culture was cooled to 20 °C before inducing protein expression with 500 µM isopropyl β-D-1-thiogalactopyranoside (IPTG). Cells were harvested 18 h post-induction by centrifugation (4,200 x g, 15 min, 4 °C) and the pellet was stored at −80 °C.

### Purification of *Staphylococcus* and *Streptococcus* FtsW: general protocol

For purification of FtsW, cells were resuspended in 50 mL lysis buffer A (50 mM HEPES pH 7.5, 0.5 M NaCl) supplemented with 1 tablet of cOmplete EDTA-free Protease Inhibitor Cocktail. Cells were lysed by passaging the resuspended cells through a cell disruptor (EmulsiFlex-C3, Avestin) at 15,000 psi three times. Cell debris was removed by centrifugation (12,000 x g, 5 min, 4 °C) and the membrane fraction was collected by ultracentrifugation of the supernatant (100,000 x g, 1 h, 4 °C). The membrane pellet was resuspended in solubilization buffer A (50 mM HEPES pH 7.5, 0.5 M NaCl, 1% n-dodecyl β-D-maltoside (DDM), 20% glycerol) using a glass dounce tissue grinder (Wheaton). The resulting mixture was stirred for 1 h at 4 °C before ultracentrifugation (100,000 x g, 1 h, 4 °C). The resulting supernatant was supplemented with 0.5 mL preequilibrated Ni-NTA resin (Qiagen) and 20 mM imidazole and stirred for 30 min at 4 °C. The sample was then loaded onto a gravity column and washed with 25 mL wash buffer A (50 mM HEPES pH 7.5, 0.5 M NaCl, 0.05% DDM, 20% glycerol) containing 20 mM imidazole and 25 mL wash buffer A containing 40 mM imidazole. The protein was then eluted in 10 mL wash buffer A containing 300 mM imidazole. The eluate was further purified by size exclusion chromatography (SEC) with a Superdex 200 10/300 GL column equilibrated in running buffer A (50 mM HEPES pH 7.5, 0.5 M NaCl, 10% glycerol, 0.05% DDM). Fractions containing the target protein were concentrated by centrifugal filtration. The absorbance at 280 nm was measured using a NanoDrop One/One Microvolume UV-Vis Spectrophotometer (ThermoFisher Scientific) and the predicted extinction coefficient (ProtParam) was used to calculate concentration.^28^ Protein samples were then aliquoted and stored at −80 °C.

### Purification of full-length *Staphylococcus* and *Streptococcus*

PBPs: general protocol. PBPs were purified via the same protocol as FtsW, above, with the following modifications. Cells were resuspended in 50 mL lysis buffer A supplemented with 1 mM PMSF. For the solubilization buffer A and wash buffer A, 10% glycerol was used instead of 20%. For SEC, a Superose 6 10/300 GL column (GE Healthcare) was used.

### Expression and purification of *S. thermophilus* ΔTMPBP2x

For expression and purification of *S. thermophilus* ΔTMPBP2x, *E. coli* BL21(DE3) containing the expression plasmid was grown in 1 L LB supplemented with carbenicillin at 37 °C with shaking until OD_600_ was 0.6. The culture was cooled to 16 °C before inducing protein expression with 500 µM IPTG. Cells were harvested 18 h post-induction by centrifugation (4,200 x g, 15 min, 4 °C) and resuspended in 25 mL lysis buffer A supplemented with 1 mM PMSF. Cell were lysed using a cell disruptor and the supernatant was collected after ultracentrifugation (100,000 x g, 30 min, 4 °C). A preequilibrated 0.5 mL Ni-NTA resin and 20 mM imidazole was added and the resulting mixture was stirred for 30 min at 4 °C. The sample was loaded onto the gravity column and washed with 25 mL lysis buffer A supplemented with 20 mM imidazole and 25 mL lysis buffer A supplemented with 40 mM imidazole. The protein was then eluted in 10 mL lysis buffer A containing 300 mM imidazole. The eluate was further purified by SEC with a Superdex 200 10/300 GL column equilibrated in running buffer A, and the fractions containing the target protein were combined, concentrated and stored at −80 °C.

### Co-Expression and purification of *S. aureus* FtsW-PBP1

The FtsW-PBP1 complex was purified via the same protocol as *S. aureus* FtsW, above, with the following modifications. Lysis, solubilization, and wash buffers contained 150 mM NaCl. For the solubilization buffer A and wash buffer A, 10% glycerol was used. After the second ultracentrifugation, the resolubilized protein was applied to 500 µL washed α-FLAG G1 affinity resin (GenScript). The resin was washed with 25 mL wash buffer A containing 0.2% DDM followed by 25 mL wash buffer A containing 0.05% DDM. The protein was eluted into 10 mL wash buffer A supplemented with 0.5 M NaCl, 0.05% DDM, and 0.2 mg/mL FLAG peptide.

### Co-Expression and purification of *P. aeruginosa* FtsW-PBP3

For expression of *P. aeruginosa* FtsW-PBP3, *E. coli* expression strain CAM333 containing pAM174 (encodes arabinose-inducible Ulp1 protease) and the expression plasmid was grown in 2 L TB supplemented with 2 mM MgCl_2_, ampicillin, and chloramphenicol at 37 °C with shaking until OD_600_ was 0.7. The culture was cooled to 20 °C before inducing protein expression with 1 mM IPTG and 0.1% arabinose. Cells were harvested 18 h post-induction by centrifugation (4,200 x g, 15 min, 4 °C). To purify FLAG-FtsW and PBP3-His_6_, the cells were resuspended in lysis buffer B (50 mM HEPES pH 7.5, 150 mM NaCl, 20 mM MgCl_2_, 0.5 M DTT) and lysed by passage through a cell disruptor (Constant Systems) at 25,000 psi twice. Membranes were collected by ultracentrifugation (100,000 x g, 1 h, 4 °C). The membrane pellets were resuspended in solubilization buffer B (20 mM HEPES pH 7.0, 0.5 M NaCl, 20% glycerol, and 1% DDM) for 1 h at 4 °C before ultracentrifugation (100,000 x g, 1 h, 4 °C). The supernatant was supplemented with 2 mM CaCl_2_ and loaded onto a homemade M1 α-Flag antibody resin. The resin was washed with 20 column volumes (CVs) of wash buffer B (20 mM HEPES pH 7.0, 0.5 M NaCl, 20% glycerol, 2 mM CaCl_2_, 0.1% DDM) and the bound protein was eluted from the column with five CVs of elution buffer (20 mM HEPES pH 7.0, 0.5 M NaCl, 20% glycerol, 0.1% DDM, 5 mM EDTA pH 8.0, and 0.2 mg/mL 3xFLAG peptide). Fractions containing the target protein were concentrated and concentration was measured via Bradford assay. Proteins were then aliquoted and stored at −80 °C.

### Detection of PGT activity via western blot

The protocol for detecting Lipid II/peptidoglycan was adapted from previously published methods.^12,17^

*S. thermophilus* FtsW: Unless otherwise stated, proteins (*St*FtsW, 0.5 µM; bPBPs, 1 µM; SgtB and *Sp*PBP1a, 0.5 µM) were added to a 1x reaction buffer (50 mM HEPES pH 7.5, 30% DMSO, 2.5 mM MgCl_2_) containing BDL (2 mM) and *E. faecalis* Lipid II (10 µM) in a total volume of 10 µL. The samples were incubated at room temperature for 5 min (SgtB and *Sp*PBP1a) or 30 min (*St*FtsW). The reaction was heat-quenched at 95 °C for 3 min and cooled to room temperature before the addition of 1 µL *E. faecalis* PBPX (100 µM) to the reaction mixture to label Lipid II/peptidoglycan with BDL. After 30 min, 11 µL 2x Laemmli sample buffer was added to quench the labeling reaction. The samples were loaded into a 4-20% gradient polyacrylamide gel (Bio-Rad) and run at 180V. After the products were transfer to the PVDF membrane (Bio-Rad), the membrane was blocked with SuperBlock TBS blocking buffer (ThermoFisher Scientific) for 30 min. For detection of biotin-labeled products, IRDye 800CW Streptavidin (LICOR Biosciences) was added at a final concentration of 1:5000 and the membrane was incubated for 1 h. The membrane was washed 3 x 10 min with 1 x TBS, and the blots were visualized using an Odyssey CLx imaging system (LI-COR Biosciences).

*S. aureus* FtsW and FtsW-PBP1*: *Sa*FtsW and bPBP stocks (50 µM) were combined 1:1 and the mixture was chilled on ice for 30 min. The proteins (*Sa*FtsW, 2.5 µM; bPBPs, 2.5 µM; SgtB, 1 µM) were then added to a 1x reaction buffer (50 mM HEPES pH 7.5, 30% DMSO, 10 mM MgCl_2_) containing BDL (3 mM), moenomycin (2 µM) and *S. aureus* Lipid II (10 µM) in a total volume of 10 µL. The samples were incubated at room temperature for 5 min (SgtB) or 1 h (*Sa*FtsW). The reaction was heat-quenched at 95 °C for 10 min. After cooling, 0.5 µL *E. faecalis* PBPX (200 µM) was added to the reaction mixture and the samples were incubated for 45 min. Reactions were quenched by the addition of 10.5 µL 2x Laemmli sample buffer. The protocol for Western blot was identical to the one used for *St*FtsW, described above.

*P. aeruginosa* FtsW-PBP3: Proteins (FtsW-PBP3, 0.5 µM; SgtB^Y181D^, 0.5 µM) were added to a 1x reaction buffer (125 mM HEPES pH 7.5, 20 mM MnCl_2_, 2.5 mM Tween 80, 200 μM cephalexin, 30% DMSO) containing *E. faecalis* Lipid II (10 µM) in a total volume of 10 μL. The samples were incubated at room temperature for 30 min. Following the incubation, reactions were heat-quenched at 95°C for 2 min. After cooling, 2 μL BDL (20 mM) and 1 μL *S.aureus* PBP4 (50 µM) were added to the reaction mixture and the samples were incubated for 1 h. Reactions was quenched by the addition of 13 μL of 2x Laemmli sample buffer and samples were loaded onto a 4-20% polyacrylamide gel. The peptidoglycan product was then transferred onto PVDF membrane and the membrane was fixed by incubating in 0.4% paraformaldehyde in phosphate buffered saline (PBS) for 30 min. Subsequently, the blot was blocked using SuperBlock buffer. The biotin-labeled products were detected by incubation with IRDye 800CW Streptavidin (1:5,000 dilution). The membrane was then washed four times with TBS with 0.5% Tween-20 (TBST), followed by one wash with PBS prior to imaging.

### LC-MS analysis of peptidoglycan crosslinking activity

This procedure was adapted from prior reports.^16,17^ *E. faecalis* Lipid II (20 µM), PGT (0.5 µM), and *S. thermophilus* PBP2x (1 µM) were incubated in a 30 µL 1x reaction buffer (50 mM HEPES pH 7.5, 10 mM MgCl_2_, 30% DMSO) at 30 °C for 1 h. The reactions were heat-quenched at 95 °C for 3 min and allowed to cool to room temperature. Polymerized material was digested by incubating the mixture at 37 °C for 3 h after the addition of 65 µL ddH_2_O and 5 µL mutanolysin (from *Streptomyces globisporus*, 4000 U/mL). To reduce the muropeptide products, 50 µL sodium borohydride (10 mg/mL) was added and the mixture was incubated at room temperature for 30 min. The pH was adjusted to ˜4 with 20% phosphoric acid and the samples were lyophilized to dryness overnight. The samples were resuspended in 20 µL ddH_2_O and analyzed by LC-MS on an Agilent 6520 QTOF operating in positive ion mode. Muropeptide products were separated on a Waters Symmetry Shield RP18 column (5 µM, 3.9 x 150 mm) with the following method: flow rate = 0.5 mL/min, 100% solvent A (H_2_O, 0.1% formic acid) for 5 min followed by a linear gradient of solvent B (acetonitrile, 0.1% formic acid) from 0 to 40% over 25 min. Molecular ions for the target muropeptide fragments were extracted from the total ion chromatogram.

### Electron microscopy of *S. aureus* cells

Overnight cultures grown in TSB supplemented with erythromycin at 30 °C were back-diluted 1:100 in fresh TSB supplemented with or without 1 mM IPTG and grown to mid-log phase at 37 °C. Cells were then fixed by adding a mixture of 1.25% formaldehyde, 2.5% glutaraldehyde, and 0.03% picric acid in 0.1 M sodium cacodylate buffer (pH 7.4). The fixed samples were imaged by electron microscopy (JEOL 1200EX-80Kv, Harvard Medical School EM Facility) as described previously.^29^

### Phase contrast microscopy of *P. aeruginosa* cells

Overnight cultures grown in LB were back-diluted 1:500 in M9 containing 0.2% casamino acids and 0.2% glucose. These cultures were then grown to an OD_600_ of 0.2 at 30°C whereupon 1 mM of IPTG was added to induce expression. The induced cells were grown for 2.5 h. Live cells were imaged using phase contrast microscopy using a Nikon Ti inverted microscope equipped with a 100x Plan Apo 1.4 Oil Ph3 DM objective, and an Andor Zyla 4.2 Plus sCMOS camera.

### Spot dilution assay of *S. pneumoniae* strains

Cultures were grown from glycerol stocks in THY supplemented with 0.2 mM ZnSO_4_ and 0.02 mM MnCl_2_ until exponential phase and normalized to an OD_600_ of 0.1. The normalized cultures were serially diluted and 5 µL of each dilution was spotted onto TSA 5%SB plates containing 0.5% fucose or 0.2 mM ZnSO_4_. Plates were imaged after overnight incubation.

## Acknowledgements

We thank M. Sjodt and A. Kruse for advice on protein purification. Funding for this work was provided by National Institutes of Health grants R01AI083365 (to T.G.B.), R01AI099144 (to T.G.B. and S.W.), R01GM076710 (to D.K. and S.W.) and F32GM123579 (to M.A.W.). A.T. is supported in part by the Funai Overseas Scholarship.

## Author Contributions

A.T., T.G.B. and S.W. conceived the project. A.T., D.K., T.G.B. and S.W. designed and coordinated the overall study. Experiments were performed by A.T., M.A.W., L.S.M., and W.L. The manuscript was written by A.T., M.A.W., T.G.B. and S.W. with input from all authors.

